# Cannabis, Connectivity, and Coming of Age: Associations between Cannabis Use and Anterior Cingulate Cortex Connectivity during the Transition to Adulthood

**DOI:** 10.1101/776138

**Authors:** Sarah D. Lichenstein, Daniel S. Shaw, Erika E. Forbes

## Abstract

**Background:** Cannabis use is common among adolescents and emerging adults and is associated with significant adverse consequences for a subset of users. Rates of use peak between the ages of 18-25, yet the neurobiological consequences for neural systems that are actively developing during this time remain poorly understood. In particular, cannabis exposure may interfere with adaptive development of white matter pathways underlying connectivity of the anterior cingulate cortex, including the cingulum and anterior thalamic radiations (ATR).

**Methods:** The current study examined the association between cannabis use during adolescence and emerging adulthood and white matter microstructure of the cingulum and ATR among 158 male subjects enrolled in the Pitt Mother & Child Project, a prospective, longitudinal study of risk and resilience among men of low socioeconomic status. Participants were recruited in infancy, completed follow-up assessments throughout childhood and adolescence, and underwent diffusion imaging at age 20 and 22.

**Results:** At age 20, moderate cannabis use across adolescence (age 12-19) was associated with higher fractional anisotropy of the cingulum and ATR, relative to both minimal and heavy adolescent use. Longitudinally, moderate and heavy extended cannabis use (age 12-21) was associated with reduced positive change in FA in both pathways from age 20 to 22, relative to minimal use.

**Conclusions:** These results suggest that precocious white matter development may be linked to increased risk for use, whereas cannabis exposure may delay white matter maturation during the transition to adulthood and potentially impact individuals’ functioning later in development.

## Introduction

Cannabis is currently the most widely used drug of abuse, with 44% of individuals in the United States reporting lifetime use (1). Rates of use are particularly high among adolescents and young adults, and likely will continue to increase based on the increasing legalization of cannabis, with approximately 20% of individuals aged 18-25 reporting cannabis use in the last month (1). Despite growing public perception of cannabis as benign (2), cannabis use can have significant deleterious consequences for some users, including substance dependence, mental health problems, and poor psychosocial attainment (3). Identifying individuals who are at the highest risk for these negative consequences could facilitate the development of targeted prevention and intervention programs to mitigate cannabis’ long-term deleterious impact. Therefore, it is imperative to elucidate the mechanisms underlying cannabis effects to predict when and for whom cannabis use is most likely to lead to negative long-term outcomes.

Widespread developmental changes in neural structure and function contribute to improvements in cognitive, affective, and social domains across adolescence (4). Various neural systems follow distinct courses of development, creating cascading points of vulnerability when environmental insults can have a disproportionately profound effect on the long-term development of different systems throughout the brain (5). Much of existing research on adolescent brain development and substance use has focused on mid-adolescence—the typical age of onset—as a time of heightened risk for use and increased vulnerability to the effects of exposure. Nonetheless, cannabis use peaks during the transition to adulthood in the early-20s—a time at which brain development, especially white matter development, remains underway—yet little research has focused on this time point. Therefore, it would be important to examine neurodevelopmental processes that may be particularly vulnerable to cannabis effects during this later period of peak use.

White matter is thought to be an important target of cannabis effects on the brain during adolescence and emerging adulthood. Comprised of myelinated axon bundles (known as white matter pathways or tracts), white matter provides the structural basis for neural signaling and undergoes significant developmental changes into early adulthood (6). Chronic cannabis exposure is associated with a downregulation of endogenous cannabinoid (CB1) receptors in the brain (7), which may interfere with normative endocannabinoid functioning and negatively impact white matter integrity via increased neuroinflammation (8-11), reduced oligodendrocyte survival (12, 13) and/or decreased myelination (14).

In particular, white matter pathways that support connectivity of the anterior cingulate cortex (ACC) may be critical targets of cannabis effects during the transition to adulthood. The ACC plays an important role in integrating cognitive, affective, and social neural networks to guide behavior (15), and has been hypothesized to function as a hub for internetwork connectivity (16, 17). Although the basic architecture of ACC connectivity remains stable from childhood, changes in the strength of various white matter pathways facilitate specialization and integration of neural networks across adolescence (17). Specifically, the cingulum and anterior thalamic radiations (ATR) are the primary white matter pathways linking the ACC with its distributed cortical and subcortical targets. The cingulum runs along the anterior-posterior axis of each hemisphere and facilitates communication between frontal, parietal, and temporal cortices as well as subcortical structures, whereas the ATR is the primary conduit for communication between the ACC and subcortical limbic structures. Each of these pathways follows a protracted developmental course, reaching estimated peaks at ages 34 and 28, respectively (16). Thus, the developmental period of greatest prevalence of problem-level cannabis use coincides with ongoing development in the structural connections that facilitate the sophisticated, circuit-based functioning of the brain.

ACC connectivity may be particularly sensitive to the effects of cannabis exposure during the transition into adulthood. Chronic cannabis users exhibit a particularly marked downregulation of CB1 receptor density in the ACC and neocortex (7), and late adolescent/young adult cannabis users exhibit altered local network organization of the cingulate cortex relative to controls (18). Congruently, cross-sectional (19-25) and longitudinal (26-29) evidence indicates altered cingulum and ATR microstructure among cannabis users. However, cross-sectional studies report both increased and decreased white matter integrity, and existing longitudinal reports are limited based on small sample sizes (maximum *N*=48 to date) and inconsistency in the age range studied. The current study builds upon previous literature by utilizing a large sample of low income, urban men, a population with particularly high rates of cannabis use. Additionally, we targeted the transition to adulthood by analyzing changes in the cingulum and ATR from age 20 to 22, allowing us to examine longitudinal associations between cannabis use and developing white matter.

Specifically, the objectives of the current study were to examine the association between cannabis use and: 1) ACC connectivity at age 20 and 2) developing ACC connectivity from age 20 to 22. We hypothesized that: 1) adolescent cannabis use (age 12-19) would predict poorer white matter integrity at age 20 (i.e. lower fractional anisotropy (FA)), and 2) extended cannabis use across adolescence and the transition to adulthood (age 12-21) would be associated with less positive change in FA from age 20 to 22 in the cingulum and ATR.

## Materials and Methods

The current sample (*n*=158) was 51.3% European American, 41.1% African American and 7.6% other races (see Table 1 for subject characteristics). Participants were characterized by low family income in early childhood, internalizing and externalizing symptoms in the normal range in early adolescence (age 10-12), and normal IQ. Less than 14% reported a non-substance-related psychiatric disorder at age 20 or 22.

**Table 1.** Sample Demographic and Clinical Characteristics. Demographic and clinical characteristics, as well as psychosocial outcome measures are presented for the full sample (N=158), and for each extended cannabis use group separately. *Note*. \**p*<.05 \*\**p*<.01. Superscript numbers in parentheses indicate which groups were significantly different from one another, based on pairwise Bonferroni-corrected post-hoc testing or pairwise χ2 tests, as applicable (1=Minimal/No Cannabis Exposure Group, 2=Moderate Cannabis Exposure Group, 3=Heavy Cannabis Exposure Group). Race was assessed by self report: participants indicated whether they identified as Asian, Black/African American, Caucasian/White, Hispanic, Mexican American, Native American, Native Hawaiian, Biracial, or Other. Subjects included in the current analyses endorsed 4 different racial classifications, black, white, biracial, or other, and data were not further reduced. Internalizing and externalizing symptoms were measured using the Child Behavior Checklist (CBCL) (Achenbach & Edelbrock, 1983), parent report, at child age 10, 11, and 12. IQ was assessed using a short form of the Wechsler Intelligence Scale for Children (WISC-III) (Wechsler, 1991). Mood Disorder includes Major Depressive Disorder (MDD; age 20 n=15, age 22 n=20), Bipolar Disorder (age 20 n=2, age 22 n=1), and dysthymia (age 20 n=3, age 22 n=5). Anxiety Disorder includes Panic Disorder (age 20 n=2, age 22 n=3), Social Phobia (age 20 n=12, age 22 n=9), Specific Phobia (age 20 n=5, age 22 n=8), Obsessive Compulsive Disorder (OCD; age 20 n=4, age 22 n=3), Post-Traumatic Stress Disorder (PTSD; age 20 n=3, age 22 n=2), and Generalized Anxiety Disorder (GAD; age 20 n=1, age 22 n=2). Educational attainment was measured on a 13-point scale, ranging from 1 – below grade 9, to 13 – completion of graduate degree.

|  |  | Full Sample (*n*=158) |  | Minimal/No Cannabis Exposure (*n*=53) |  | Moderate Cannabis Exposure (*n*=52) |  | Heavy Cannabis Exposure (*n*=53) |  | Group Comparison |  |
| --- | --- | --- | --- | --- | --- | --- | --- | --- | --- | --- | --- |
| **Race** | | *N* | % | *N* | % | *N* | % | *N* | % | $\chi^2$ | *p* |
|  | White | 81 | 51.3 | 32 | 60.4 | 24 | 46.2 | 25 | 47.2 | 5.87 | 0.437 |
|  | Black | 65 | 41.1 | 16 | 30.2 | 24 | 46.2 | 25 | 47.2 |  |  |
|  | Biracial | 8 | 5.1 | 3 | 5.7 | 2 | 3.8 | 3 | 5.7 |  |  |
|  | Other | 4 | 2.53 | 2 | 3.8 | 2 | 3.8 | 0 | 0 |  |  |
| **SES** |  | *M* | *SD* | *M* | *SD* | *M* | *SD* | *M* | *SD* | *F* | *p* |
| Family Income (per month)<br>(Mean first 3 assessments) |  | 1208.18 | 669.9 | 1236.88 | 594.24 | 1409.28<sup>(3)</sup> | 768.12 | 982.16<sup>(2)</sup> | 574.26 | 5.74 | 0.004** |
| Neighborhood Risk Score (Mean<br>first 3 assessments) |  | 0.37 | 1.12 | 0.03<sup>(3)</sup> | 0.81 | 0.45 | 1.25 | 0.63<sup>(1)</sup> | 1.2 | 4.06 | 0.019* |
| **Childhood Clinical and Cognitive Assessments** |  |  |  |  |  |  |  |  |  |  |  |
| Internalizing Symptoms (Mean |  | 5.28 | 4.99 | 5.38 | 5.14 | 4.95 | 4.82 | 5.49 | 5.07 | 0.55 | 0.581 |
| Externalizing Symptoms (Mean |  | 8.65 | 6.85 | 7.98 | 6.51 | 7.77 | 6.22 | 10.09 | 7.55 | 0.15 | 0.865 |
| IQ (Age 11) |  | 96.11 | 18.39 | 97.89 | 20.82 | 96.61 | 19.94 | 93.86 | 13.94 | 1.68 | 0.19 |
| **Age 20 Psychopathology** | | *N* | % | *N* | % | *N* | % | *N* | % | $\chi^2$ | *p* |
| Mood Disorder |  | 18 | 11.39 | 5 | 3.16 | 4 | 2.53 | 9 | 5.7 | 2.49 | 0.287 |
| Anxiety Disorder |  | 21 | 13.29 | 6 | 3.8 | 5 | 3.16 | 10 | 6.33 | 2.17 | 0.339 |
| Psychosis |  | 3 | 1.9 | 2 | 1.27 | 0 | 0 | 1 | 0.63 | 2.05 | 0.358 |
| Antisocial Personality Disorder |  | 12 | 7.6 | 0<sup>(3)</sup> | 0 | 0<sup>(3)</sup> | 0 | 12<sup>(1,2)</sup> | 22.64 | 25.5 | <.001** |
| **Age 22 Psychopathology** |  |  |  |  |  |  |  |  |  |  |  |
| Mood Disorder |  | 22 | 13.9 | 8 | 15.1 | 7 | 13.5 | 7 | 13.2 | 0.106 | 0.948 |
| Anxiety Disorder |  | 20 | 12.7 | 3 | 5.7 | 5 | 9.6 | 12 | 22.6 | 7.69 | 0.021 |
| Psychosis |  | 3 | 1.9 | 1 | 1.9 | 1 | 1.9 | 1 | 1.9 | 0 | 1 |
| Antisocial Personality Disorder |  | 17 | 10.8 | 1<sup>(3)</sup> | 1.9 | 0<sup>(3)</sup> | 0 | 16<sup>(1,2)</sup> | 30.2 | 31.82 | <.001** |
| **Psychosocial Outcome Measures** |  |  |  |  |  |  |  |  |  |  |  |
| Completed HS/GED (or less)<br>(Age 22) |  | 76 | 48.1 | 22<sup>(3)</sup> | 41.5 | 15<sup>(3)</sup> | 28.8 | 39<sup>(1,2)</sup> | 73.6 | 22.43 | <.001** |
| Currently Employed (Age 23) |  | 47 | 29.7 | 12 | 22.6 | 17 | 32.7 | 18 | 34 | 3.19 | 0.202 |

### Pitt Mother and Child Project (PMCP; Shaw, Gilliom, Ingoldsby, & Nagin, 2003)

Participants were part of the Pitt Mother & Child Project, a longitudinal study of risk and resilience among men from families of low socioeconomic status (SES). A total of 310 mother-son dyads were recruited from Women, Infant, and Children (WIC) Nutritional Supplement centers in the Pittsburgh metropolitan area when subjects were 6-17 months old, and they have been followed throughout childhood, adolescence, and into young adulthood (in-person home, lab, and/or internet/phone assessments at ages 1.5, 2, 3.5, 5, 5.5, 6, 8, 10, 11, 12, 15, 16, 17, 18, 20, 21, 22, and 23) (30). All procedures were approved by the Institutional Review Board at the University of Pittsburgh, and all experiments were performed in accordance with relevant guidelines and regulations.

Participants were excluded from the MRI portion of the study if they endorsed any standard MRI contraindications. Out of the full sample (*N*=310), *n*=186 completed an MRI scan at age 20 (*n*=31 declined, *n*=17 could not be contacted, *n*=10 were incarcerated, *n*=5 lived out of the area, *n*=5 were in the military, *n*=1 was deceased, and n=56 endorsed contraindications to MRI). Of those completing the scan at age 20, 28 did not complete a second scan, resulting in a subset of *n=*158 male PMCP participants for whom good-quality DTI data were acquired at both age 20 and 22.

## Measures

### Cannabis use

Lifetime cannabis use was assessed with the Lifetime History of Drug Use and Drug Consumption (LHDU; Day et al., 2008; Skinner, 1982) semi-structured interview at age 20 and 22. Participants who endorsed a positive lifetime history of cannabis use (at least 3 times in one year) reported their age of cannabis use onset, annual frequency of use, and their greatest use in one day for each year since their first use.

Adolescent cannabis use was quantified by calculating the sum of participants’ average days of use/month at each time point from age 12-19. Because participants were scanned around their 20^th^ birthday, age 19 represents the year preceding their baseline DTI scan. Because these data were not normally distributed and contained a significant proportion of zero values, the sample was divided into terciles based on total frequency of use from age 12-19, which consisted of a minimal adolescent use group (*n*=56; ≤1 days/month), a moderate adolescent use group (*n*=49; ∼weekly use), and a heavy adolescent use group (*n*=53; multiple uses/week).

Extended cannabis use across adolescence and the transition to adulthood was measured by calculating the sum of participants’ average days of use/month at each time point from age 12-21. Because participants were scanned around their 22^nd^ birthday, age 21 represents the year preceding their follow-up DTI scan. Again, participants were split into terciles, with *n*=53 participants assigned to a minimal extended use group (<1x/week), *n*=52 assigned to a moderate extended use group (<1.5 times/week), and *n*=53 assigned to a heavy extended use group (multiple uses/week).

### Alcohol use

Alcohol use was assessed with the Lifetime History of Alcohol Use and Alcohol Consumption semi-structured interview (33) at age 20. Alcohol use frequency and number of drinks were multiplied to obtain a measure of overall quantity of alcohol exposure for each year since first use. Alcohol exposure for each year from age 13-19 was summed to create a measure of lifetime alcohol exposure, and a log transformation was used to account for a positive skew in the data.

### Race

Self-report of race at age 20 was used. Participants reported whether they identified as Asian, Black/African American, Caucasian/White, Hispanic, Mexican American, Native American, Native Hawaiian, Biracial, or Other.

### Head motion

Mean head displacement was calculated for each participant for each DTI scan.

### Diffusion Tensor Imaging (DTI)

Participants underwent DTI scanning at age 20 and 22 on a 3T Siemens Tim Trio scanner at the University of Pittsburgh MR Research Center. Diffusion-sensitizing gradient encoding was applied in 61 uniform angular directions with a diffusion weighting of b=1000 s/mm^2^. Preprocessing was carried out using the Oxford Centre for Functional MRI of the Brain (FMRIB) Software Library (FSL) (34) using tract-based spatial statistics (TBSS) (35). The Johns Hopkins University White Matter Tractography Atlas (36) was used to identify the right and left cingulum (cingulate gyrus) and ATR as regions of interests (ROIs), and mean FA, MD, AD, and RD values for each ROI were extracted to SPSS for further analysis. Details on the DTI methods are presented in the Supplementary Materials.

### Data Analysis

Analysis of covariance (ANCOVA) was used to determine whether microstructure of the cingulum and ATR at age 20 differed between adolescent cannabis use groups. Separate models were constructed for FA, MD, AD and RD of the cingulum and ATR. Alcohol use, tobacco use, IQ, SES, and psychopathology were considered as covariates based on reported associations with measures of white matter integrity. To determine the appropriate covariates to include in the final models, the Akaike Information criterion (AIC) was used to compare candidate models and any covariate that substantially improved the model fit (difference in AIC ≥ 2) was included in the final analyses (see Table S1). Based on these analyses, models for age 20 FA included race as a covariate. Additionally, although it did not significantly improve model fit, models including alcohol use were also tested (Table S3-S4) based on evidence that accounting for alcohol use may attenuate cannabis associations with brain structure (32) (measures for assessing covariates not included in the final analyses are presented in Supplementary Materials). For all analyses, hemisphere and head motion were also included as covariates (31), subject ID number was included as a random effect variable, and Bonferroni-corrected post-hoc pairwise tests were used to probe significant main effects.

An exploratory whole-brain analysis was also performed using the *randomise* tool in FSL (37) to assess effects of adolescent cannabis exposure throughout the brain. A voxel-based FWE-corrected significance threshold of *p*<.01 was used to evaluate results. Any additional regions identified in which age 20 microstructure differed significantly between adolescent use groups were also to be included in subsequent longitudinal analyses.

Finally, ANCOVAs were computed to estimate whether change in microstructure of the cingulum or ATR from age 20-22 (i.e., difference score for FA from age 20 to 22) varied between extended cannabis use groups.

## Results

### Sample Characteristics

#### Cannabis use

Consistent with the high prevalence of use in this population (compared with 52.7% in a nationally representative sample (1)), 79% of participants (*n*=124) reported a lifetime history of cannabis use. No participants reported regular use prior to age 12 (see Table 2 for details on cannabis use).

**Table 2.**
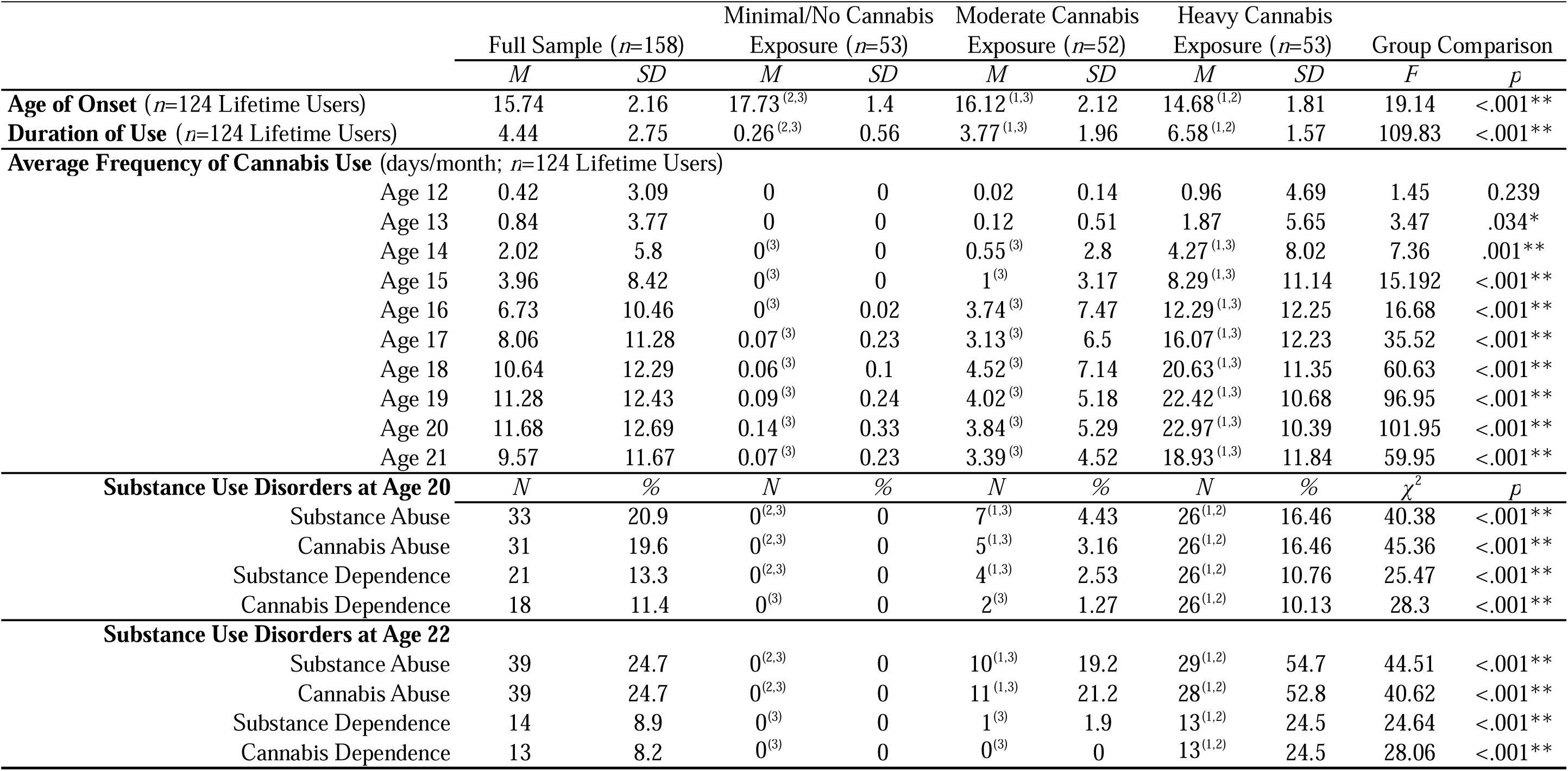
Cannabis Use Characteristics. Details regarding cannabis use onset, duration, annual frequency of use, and substance use disorder diagnoses are presented for the full sample (N=158), and each extended cannabis use group separately. *Note*. \**p*<.05 \*\**p*<.01. Superscript numbers in parentheses indicate which groups were significantly different from one another, based on pairwise Bonferroni-corrected post-hoc testing or pairwise χ2 tests, as applicable (1=Minimal/No Cannabis Exposure Group, 2=Moderate Cannabis Exposure Group, 3=Heavy Cannabis Exposure Group). Duration of use reflects the number of years (from age 12-21) that participants reported cannabis use frequency ≥ 1x/week. Substance use disorder diagnoses determined based on the Structured Clinical Interview for DSM-IV (SCID), administered at age 20 and 22 study visits. Substance Abuse: Includes Cannabis Abuse (age 20 n=31, age 22 n=39), Sedative Abuse (age 20 n=0, age 22 n=4), Stimulant Abuse (age 20 n=0, age 22 n=1), Opioid Abuse (age 20 n=2, age 22 n=3), Cocaine Abuse (age 20 n=2, age 22 n=2), Hallucinogen/PCP Abuse (age 20 n=2, age 22 n=2), and Other Substance Abuse (age 20 n=1, age 22 n=1). Substance Dependence: Includes Cannabis Dependence (age 20 n=18, age 22 n=13), Sedative Dependence (age 20 n=1, age 22 n=2), Opioid Dependence (age 20 n=1, age 22 n=3), Cocaine Dependence (age 20 n=1, age 22 n=22), Hallucinogen/PCP Dependence (age 20 n=0, age 22 n=1), and Other Substance Dependence (age 20 n=1, age 22 n=0).

#### Alcohol and other illicit substance use

Ninety-six percent of participants (*n*=151) reported a lifetime history of alcohol use. Cumulative alcohol exposure was higher among those with higher rates of cannabis use. Less than 15% of participants reported lifetime use of illicit drug other than cannabis (see Table S2 for additional information on alcohol and other substance use).

### Cross-sectional Association between Adolescent Cannabis Use and ACC Connectivity at Age 20

#### Cingulum

##### FA

Effects on cingulum FA at age 20 were observed for adolescent cannabis use group (Table 3; Figure 1). Contrary to our hypothesis, the moderate use group displayed higher FA than both other groups. None of the pairwise differences among groups met the Bonferroni-corrected threshold. However, when alcohol use was included as a covariate (Table S3), the cannabis group effect remained and pairwise comparisons indicated greater cingulum FA in the moderate versus minimal use group.

**Table 3.** Cross-sectional Association between Adolescent Cannabis Use and ACC Connectivity at Age 20. Significant effects on FA and MD of the cingulum and anterior thalamic radiations were observed for adolescent cannabis use group and hemisphere. Additionally, significant effects of head motion and race, as well as the interaction between adolescent cannabis use group and race, were also observed for cingulum FA. *Note*. Each quadrant represents one ANCOVA and significant effects (*p*<.05) are bolded. FA=fractional anisotropy, ATR=anterior thalamic radiations, *df*=degrees of freedom.

|  | Age 20 FA |  |  |  |  |  |
| --- | --- | --- | --- | --- | --- | --- |
|  | Cingulum |  |  | ATR |  |  |
|  | *F* | *df* | *p* | *F* | *df* | *p* |
| Adolescent Cannabis Use Group | **2.96** | **2** | **0.05** | **6.29** | **2** | **0.00** |
| Hemisphere | **12.67** | **1** | **0.00** | **15.75** | **1** | **0.00** |
| Head Motion | **4.24** | **1** | **0.04** | 0.46 | 1 | 0.50 |
| Race | **4.85** | **3** | **0.00** | 0.12 | 3 | 0.95 |
| Cannabis Group*Race | **2.82** | **5** | **0.02** | - | - | - |
|  | Age 20 MD |  |  |  |  |  |
|  | Cingulum |  |  | ATR |  |  |
|  | *F* | *df* | *p* | *F* | *df* | *p* |
| Adolescent Cannabis Use Group | **7.39** | **2** | **0.00** | **3.48** | **2** | **0.03** |
| Hemisphere | **12.72** | **1** | **0.00** | **8.19** | **1** | **0.00** |
| Head Motion | 0.90 | 1 | 0.35 | 0.09 | 1 | 0.76 |

**Figure 1.**
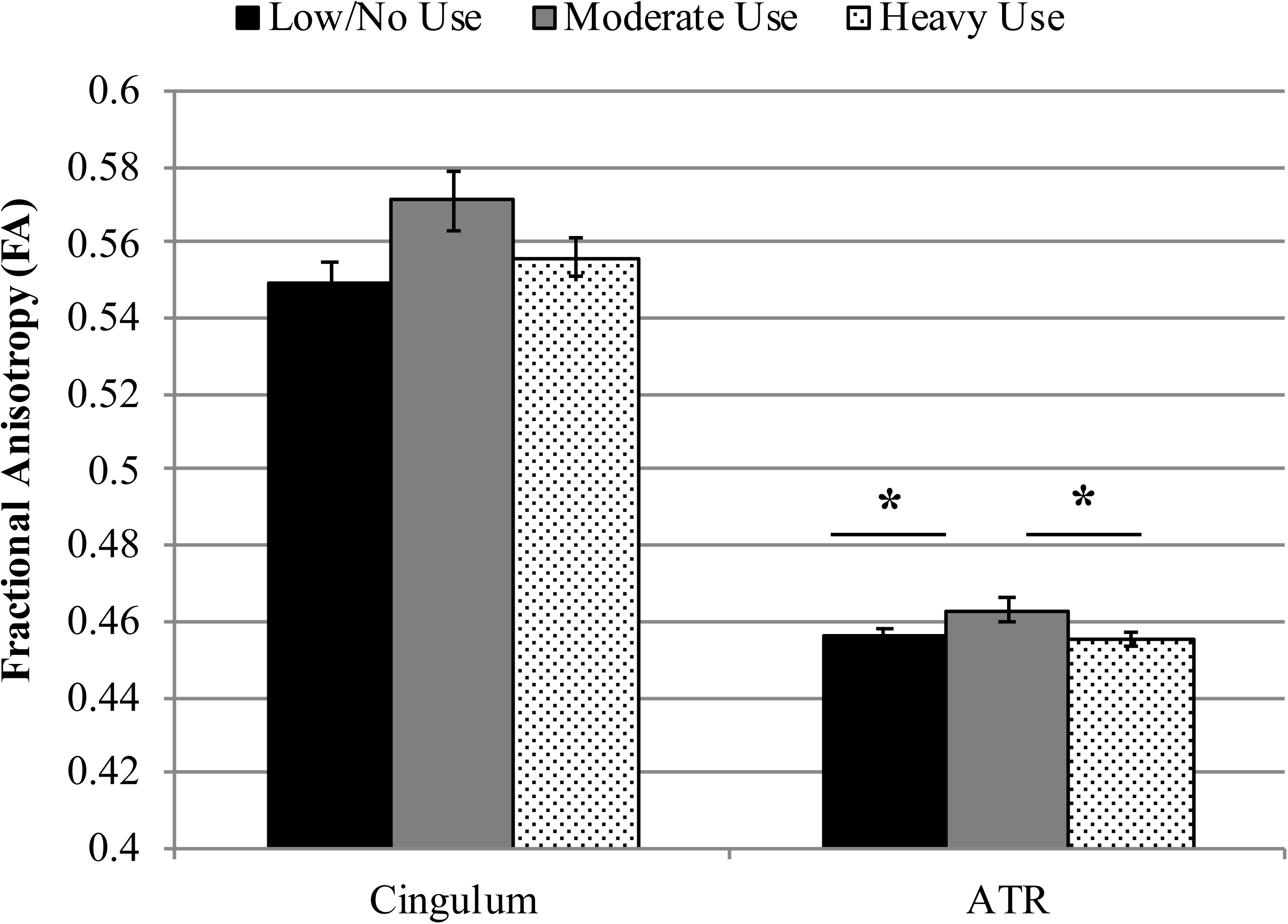
Main effect of adolescent cannabis exposure on cingulum and ATR FA at age 20. Significant main effects of adolescent cannabis exposure group were observed for both the cingulum and anterior thalamic radiations. For the anterior thalamic radiations, Bonferroni-corrected pairwise post-hoc tests demonstrated that the moderate use and low or no use groups differed (*p*_corrected_<.05), as did the moderate and heavy use groups (*p*_corrected_<.01).

An adolescent cannabis use group by race interaction was also found (Figure 2), such that among African American participants there was no effect of cannabis exposure (*F*=.58, *ns*), whereas cannabis use group was significantly associated with FA among European Americans (*F*=11.01, *p*<.001). Within-group associations were not examined for biracial subjects (*n*=8) and subjects of other races (*n*=4) because of insufficient sample sizes.

**Figure 2.**
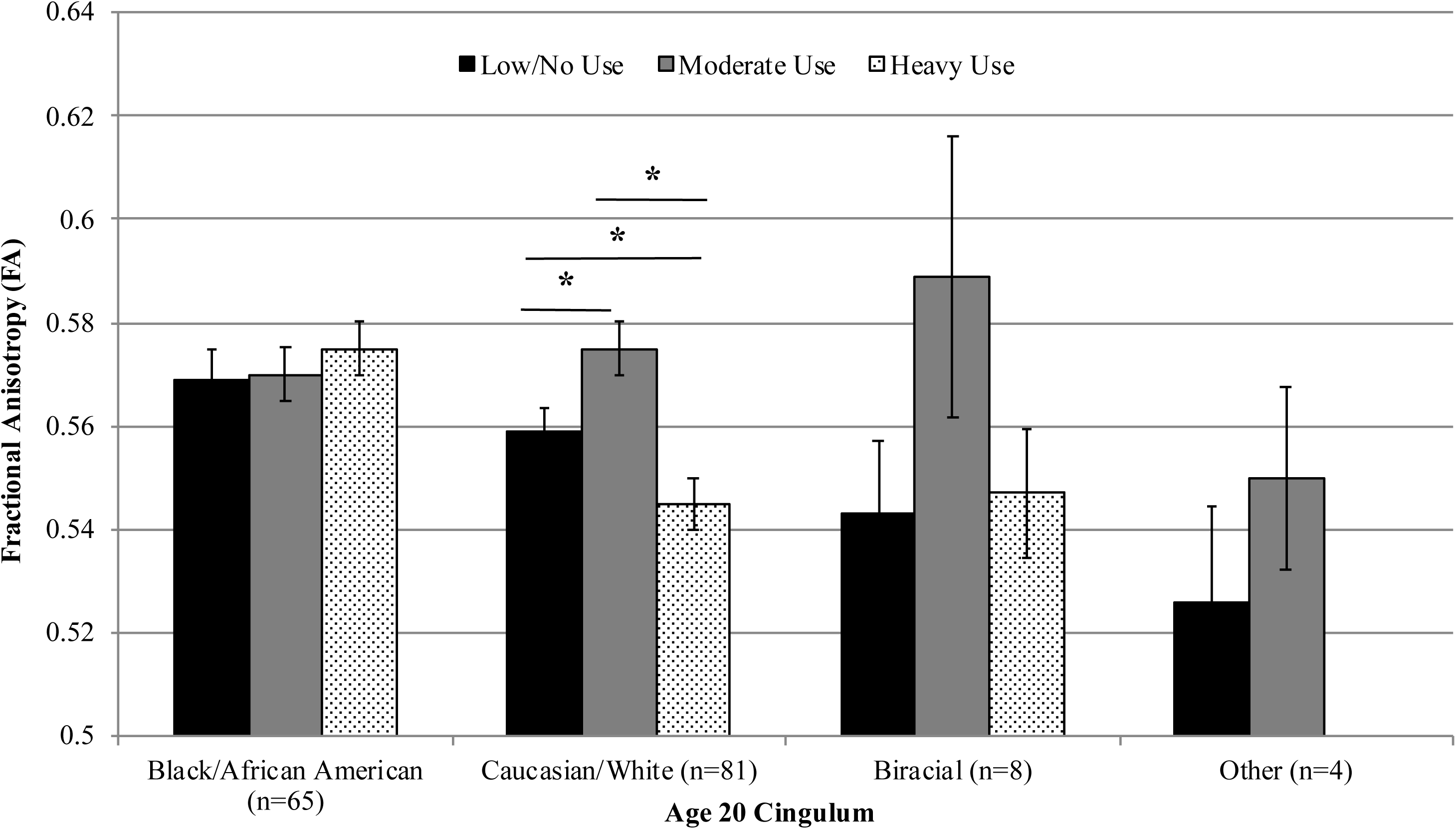
Interactive effect of cannabis exposure and participant race on cingulum FA at age 20. A significant cannabis exposure by race interaction was found, such that among African American participants there was no significant effect of adolescent cannabis exposure whereas among European American participants, adolescent cannabis exposure had a significant main effect on cingulum FA (*F*=11.01, *p*=.000). Among European American participants, moderate cannabis use was linked to greater FA relative to low or no cannabis use (*p*_corrected_<.05), whereas heavy adolescent cannabis use was associated with lower FA relative to both the low or no use group (*p*_corrected_<.05) and the moderate use group (*p*_corrected_<.001). Within group associations were not examined for biracial subjects (n=8) and subjects of other races (n=4) due to insufficient sample sizes.

##### MD

For cingulum MD, effects were again observed for adolescent cannabis use group (Table 3). The moderate use group had greater MD than the heavy use group (*p*_corrected_<.001). The effect of adolescent cannabis use group and post-hoc results remained significant when alcohol use was included as a covariate (Table S3).

#### ATR

##### FA

Significant effects on ATR FA were also observed for adolescent cannabis use group (Table 3; Figure 1). Similar to the pattern observed for cingulum FA, the moderate adolescent use group displayed higher FA than both the minimal and the heavy adolescent use groups (*p*_corrected_’s<.05). The adolescent cannabis use group effect and post hoc results remained significant after controlling for alcohol (Table S3).

##### MD

Effects were observed for adolescent cannabis use group (Table 3). The moderate use group had higher MD than the heavy use group (*p*_corrected_<.05). This effect was no longer significant when alcohol was included, although the effect of alcohol exposure was not significant (Table S3).

##### Whole-brain results

The whole-brain analysis did not reveal any clusters throughout the white matter skeleton in which white matter microstructure differed significantly between adolescent cannabis use groups. Therefore, subsequent analyses of longitudinal cannabis effects do not include additional ROIs.

### Longitudinal Association between Extended Cannabis Use and Developing ACC Connectivity from age 20-22

#### Cingulum

##### FA change from age 20 to 22

All extended cannabis use groups displayed increased FA from age 20 to 22, but FA change differed by group across these 2 years (Table 4; Figure 3). The 2-year increase in FA was significantly larger for the minimal relative to the moderate extended use group (*p*_corrected_<.05). The effect remained significant when controlling for alcohol (Table S4).

**Table 4.**
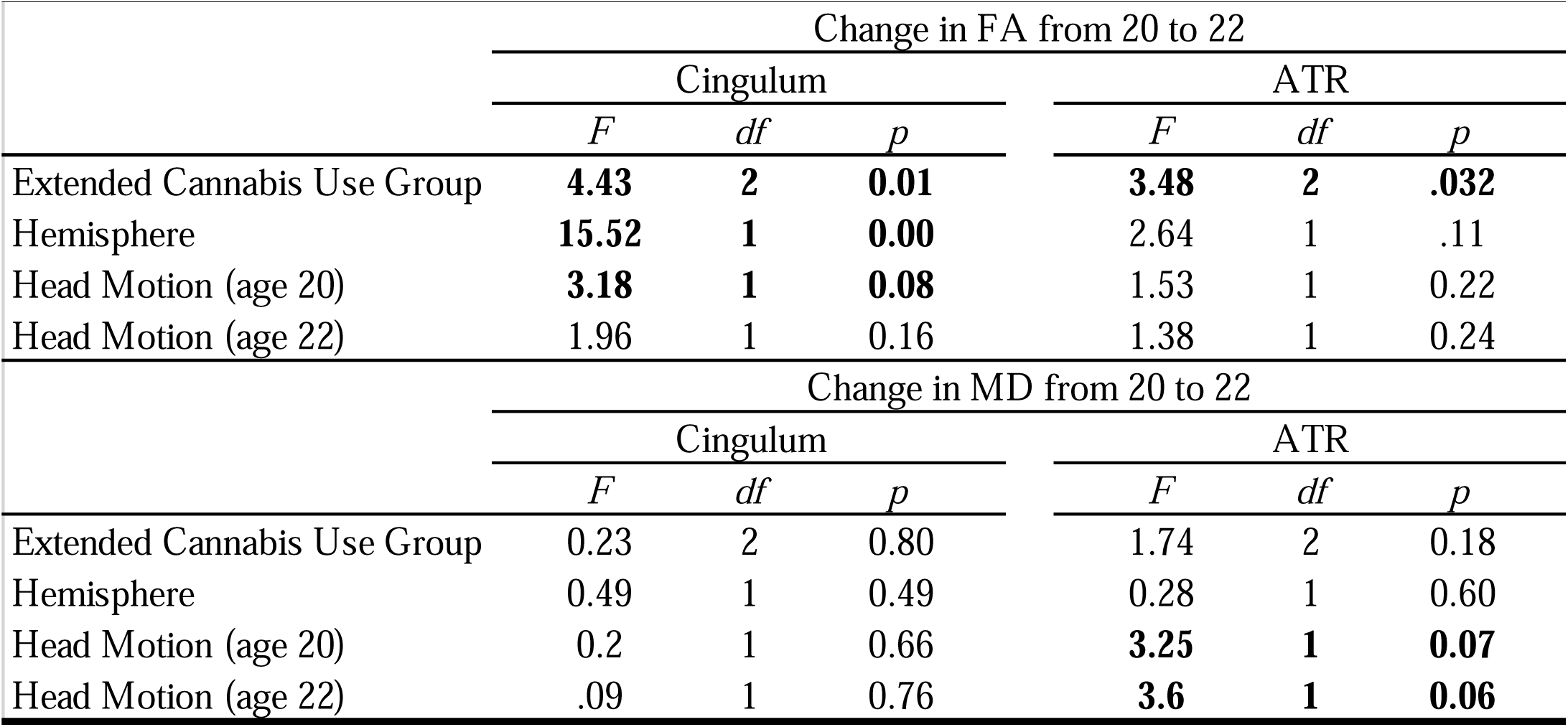
Longitudinal Association between Extended Cannabis Use and Developing ACC Connectivity from age 20-22. Significant effects on change in FA from 20 to 22 were observed for extended cannabis use group in the cingulum and anterior thalamic radiations. Additionally, significant effects of hemisphere and head motion at age 20 were observed for change in cingulum FA from 20 to 22, and significant effects of head motion at age 20 and 22 were observed for change in ATR MD from 20 to 22. Note. Each quadrant represents one ANCOVA and significant effects (p<.05) are bolded. Change in FA from 20 to 22 represents the difference score for FA from age 20 to 22, and change in MD from 20 to 22 represents the difference score for MD from age 20 to 22. FA=fractional anisotropy, ATR=anterior thalamic radiations, df=degrees of freedom.

**Figure 3.**
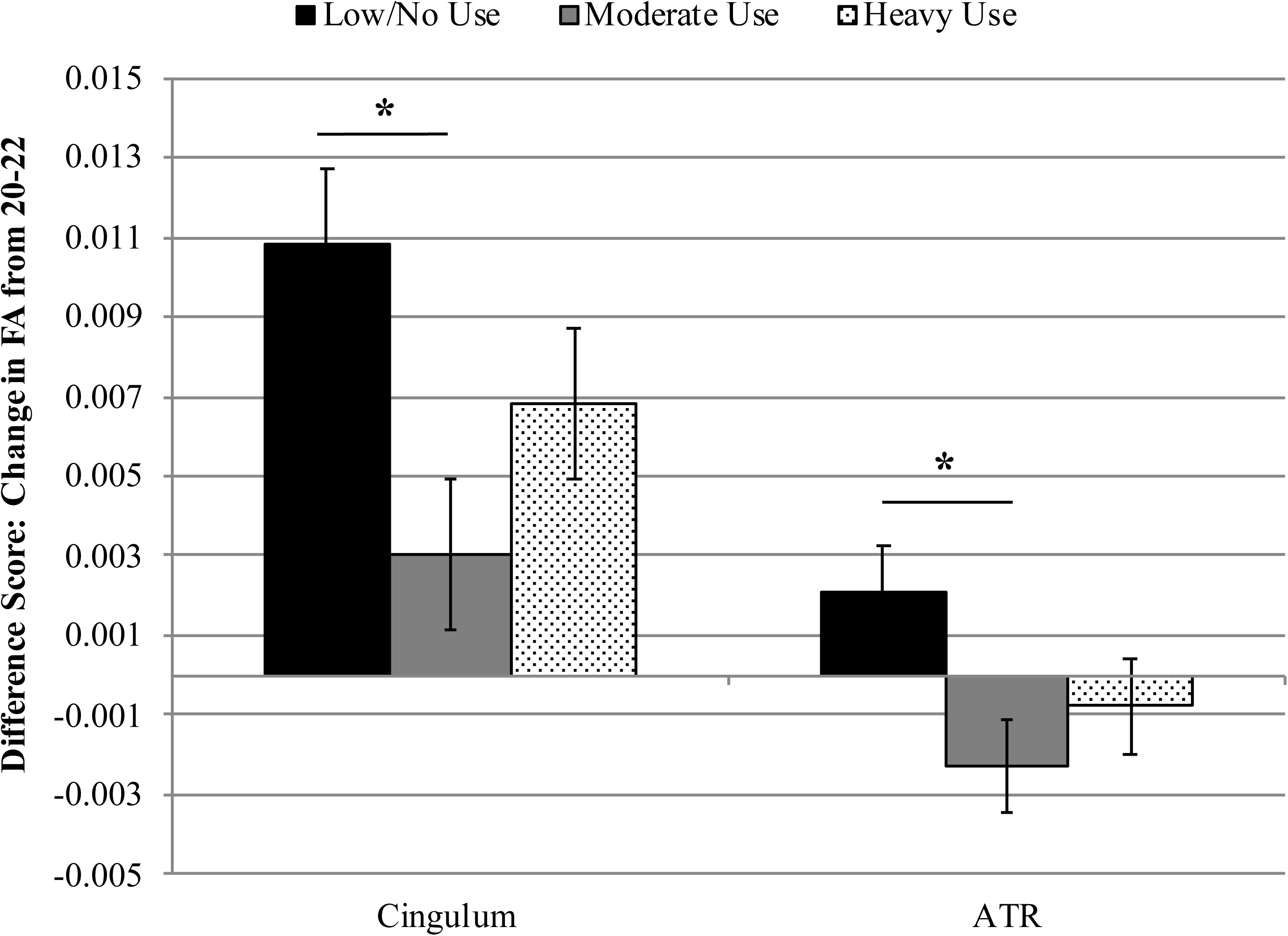
Longitudinal effects of cannabis exposure on FA development from 20 to 22. Significant effects of extended cannabis exposure and hemisphere were observed for change in cingulum FA across these 2 years. The main effect of extended cannabis exposure on change in ATR FA was also statistically significant. Bonferroni-corrected post-hoc tests revealed that the difference between the minimal use and moderate extended cannabis use groups was statistically significant for change in both cingulum and ATR FA (*p*_corrected_<.05).

##### MD change from 20 to 22

No effect of cannabis use group was observed.

#### ATR

##### FA change from age 20 to 22

The ANCOVA revealed a significant effect of extended cannabis use group for change in ATR FA (Table 4; Figure 3). The minimal use group displayed an increase in FA over the 2 years, whereas both other groups displayed a decrease (*p*_corrected_<.05). This difference was not significant after controlling for alcohol, although the effect of alcohol exposure was non-significant (Table S4).

##### MD change from age 20 to 22

No effect of cannabis use group was observed.

## Discussion

The current study aimed to examine the effects of cannabis exposure on developing ACC connectivity during the transition to adulthood. Contrary to our expectations, moderate, but not heavy, adolescent cannabis users displayed higher FA and MD in the cingulum and ATR relative to minimal users at age 20, even after controlling for alcohol exposure. In contrast, our longitudinal results supported our hypothesis that cannabis exposure is associated with reduced white matter maturation – attenuated increase or decreased FA – from age 20 to 22. Among minimal extended cannabis users, FA of the cingulum and ATR increased across this 2-year period – consistent with normative development of these pathways – but this increase in white matter integrity was reduced or even reversed among moderate and heavy cannabis extended users. These results align with existing longitudinal studies (26-29), and collectively provide strong evidence that cannabis exposure during adolescence and the transition to adulthood is associated with diminished white matter maturation of the cingulum and ATR.

Taken together, the results of our cross-sectional and longitudinal analyses highlight the need to distinguish premorbid neural characteristics associated with risk for use from the neurobiological effects of cannabis exposure. However, a variety of different patterns of aberrant white matter development may contribute to risk for psychopathology (38), and both delayed (39, 40) and accelerated (41) patterns of white matter development have been identified among individuals at high familial risk for substance use. There are multiple possible paths to substance use, as well as multiple outcomes resulting from use, and both early-developing and late-developing white matter microstructure may each contribute differently to increased propensity for substance use and the development of related problems.

One possible interpretation is that accelerated white matter development could be a characteristic of those who are on a steeper developmental trajectory, which may be linked to heightened risk for substance use. Of particular relevance to the current sample, Belsky’s fast-life theory of socialization based on evolutional models proposes that familial psychosocial stress leads individuals to mature more rapidly and reproduce earlier to improve their reproductive fitness within insecure environments (42). Congruently, emerging animal and human literature suggests that early adversity may accelerate the development of cortical-subcortical connectivity (43), and early pubertal maturation has been linked to more advanced white matter development in late adolescence (44). As participants in the current study were recruited based on low SES, this sample is characterized by high rates of neighborhood impoverishment, low income, and maternal depression, among other sources of childhood adversity (45), which could potentially lead to a compensatory acceleration in white matter development. In turn, precocious white matter development may contribute to earlier autonomy, exploration, and socialization (41). Indeed, prior research has reported higher ATR FA to be linked to heightened risky behavior among adolescents, based on both self-report (46) and behavioral measures (47).

Collectively, our findings of higher FA among moderate cannabis users at age 20 coupled with decreased FA in users across 2 years suggest a pattern of white matter development in which a subset of participants are characterized by higher FA prior to the onset of cannabis use (potentially attributable to early adversity) that may increase their liability to experiment with drugs, followed by reduced white matter maturation with extended cannabis use. We have illustrated this theoretical model in Figure 4. Accordingly, higher FA may represent a marker of risk, whereas cannabis exposure is associated with poorer white matter integrity over time. Although speculative, the current model provides promising avenues for future research to disentangle neural risk factors from cannabis effects on the developing brain.

**Figure 4.**
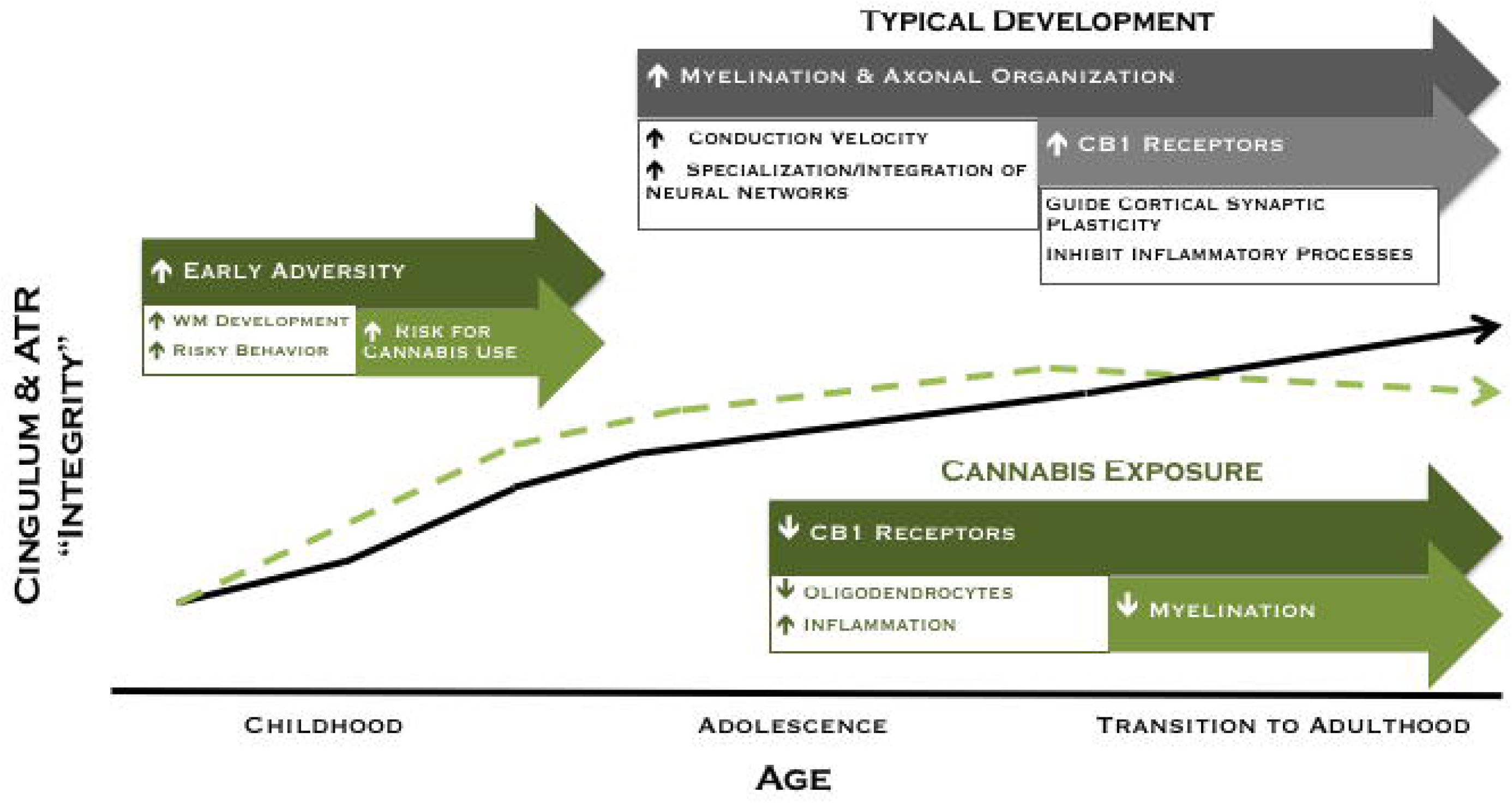
Revised theoretical model of developing ACC connectivity among individuals with and without meaningful cannabis exposure. In typical development (black line), increased myelination and axonal organization, as well as increased CB1 receptor expression are postulated to give rise to increased white matter integrity across adolescence and into adulthood. Exposure to adversity early in development may lead to a compensatory acceleration of white matter development, which may increase risky behavior and risk for cannabis use among a subset of individuals (green dashed line). Conversely, cannabis exposure is associated with reduced white matter maturation of the cingulum and ATR during the transition to adulthood, an effect that may be mediated by a downregulation of CB1 receptor expression and/or direct effects on oligodendrocyte survival and myelination.

Cannabis effects on white matter microstructure varied substantially between the moderate and heavy cannabis use groups, suggesting an effect of dose/use characteristics. At age 20, moderate adolescent users displayed the highest FA in both pathways, and the moderate extended cannabis use group was characterized by a more substantial reduction in white matter development from age 20 to 22 relative to the heavy extended use group. To our knowledge, no previous studies have compared white matter development between cannabis users with different levels of use. However, to speculate about what may be driving this pattern, the heavy users could be further along the cannabis exposure trajectory of white matter development illustrated in Figure 4. Indeed, the heavy extended use group initiated cannabis use earlier (mean age 14.7 vs. 16.1) and used more frequently than the moderate extended use group. This pattern of results is also interesting in light of previous findings from the same sample that an escalating trajectory of cannabis use across adolescence was associated with altered functional connectivity, relative to both stable low and stable high use trajectories (48). Collectively, these studies highlight the importance of considering cannabis use characteristics – including dose, timing, and trajectory – to better characterize the effects of cannabis exposure on the developing brain.

The time course of cannabis effects on the brain remains poorly understood, although data on CB1 receptor changes with cannabis use are informative. Chronic cannabis use has been linked to a downregulation of CB1 receptors (7). However, this finding was based on a case-control study including daily cannabis smokers who had been using for a mean of 12 years (7). Therefore, it is unclear whether this effect occurs quickly and is then sustained, or if it occurs gradually over the course of many years of exposure. Nonetheless, follow-up data demonstrated that receptor levels normalized after ∼4 weeks of abstinence (7), suggesting that the downregulation in receptor expression takes place on a timescale of weeks to months, not gradually over years. Therefore, cannabis effects may plateau with protracted use, which could be reflected in the current pattern of results, such that cannabis effects on change in white matter microstructure may have been more robust among moderate cannabis users because the heavy users are at a later point on the trajectory when cannabis effects have begun to plateau.

We found that adolescent cannabis use group interacted with race, such that among African American participants there was no effect of adolescent cannabis exposure, whereas adolescent cannabis exposure had a significant effect on cingulum FA at age 20 among European Americans. A paucity of research has examined whether white matter microstructure differs by race, although smaller white matter volume has been noted among Asian and African American participants relative to European Americans (49). African Americans in the current sample were characterized by lower family income and higher neighborhood risk in early childhood relative to European Americans (*p*s<.001). Because chronic stress has pervasive effects on the endocannabinoid system, including widespread downregulation of CB1 receptors (8), African American participants may be relatively less vulnerable to cannabis effects because of endocannabinoid system alterations related to elevated chronic stress, including discrimination (51, 52).

Although the current study has many strengths, including prospective, longitudinal data on adolescent/emerging adult cannabis use and white matter microstructure in a large sample of high-risk young men, there are also several limitations. We investigated a population at high risk for both cannabis use and its adverse consequences (53), but our results may not be generalizable to women, individuals of higher SES, or participants from suburban or rural communities. Additionally, the current study relies on retrospective self-reports of cannabis use. Future studies would benefit from estimates from multiple sources, prospective measurement of use, and details on cannabinoid composition (55, 56). Finally, although TBSS is relatively robust to the effects of fiber anatomy, metrics derived from the tensor model are highly susceptible to distortion from complex fiber geometry (57). Optimally, as with the ongoing ABCD study, we will learn about altered pattern and pace of white matter maturation in cannabis users through investigations that use prospective, longitudinal designs, with detailed measurements of individuals’ social, cultural, and developmental context.

## Conclusions

Cannabis use is common during adolescence and the transition to adulthood. Although often considered benign, cannabis use has been associated with a wide array of negative outcomes that can have profound impacts on individuals’ long-term trajectory of achievement, health, and well-being. However, the neurobiological mechanisms that underlie the deleterious effects of cannabis exposure, especially at vulnerable developmental periods and in high-risk populations, remain poorly understood. The current study used longitudinal DTI data to demonstrate that cannabis use is associated with lesser white matter maturation of the cingulum and ATR from age 20 to 22. These results have important implications for understanding cannabis effects on brain structure and function and informing public perceptions about the risks of cannabis use. Elucidating the neural basis of cannabis effects can facilitate the development of targeted prevention and intervention strategies to foster positive development among individuals at highest risk for cannabis use and poor psychosocial adjustment.

## Acknowledgments

This work was supported by a grant from the National Institutes of Health to Drs. Daniel S. Shaw and Erika E. Forbes (R01 DA026222). Manuscript preparation was supported in part by a National Science Foundation Graduate Research Fellowship (GRFP DGE-1247842) and National Institute of Drug Abuse Institutional Research Training Grant (T32 DA022975) awarded to Sarah Lichenstein.

The authors thank the young men and mothers who have participated in the Pitt Mother and Child Project, Dr. Timothy Verstynen for his assistance with the diffusion imaging analysis, and the staff members who assisted with data collection.

