## Supplementary Material for "Cannabis, Connectivity, and Coming of Age: Associations between Cannabis Use and Anterior Cingulate Cortex Connectivity during the Transition to Adulthood"

**Materials and Methods**

**Measures**

***Tobacco use.*** Each participant was classified based on whether or not they reported daily use of tobacco during the last year on The Alcohol and Drug Consumption Questionnaire (ADCQ; Cahalan, Cisin, & Crossley) at age 20.

***IQ.*** Prorated full Scale IQ (FSIQ) scores were derived from participants’ performance on a short form of the Wechsler Intelligence Scale for Children (WISC-III; Wechsler, 1991) at age 11.

***Socioeconomic status (SES).*** The current study used a composite measure including both familial income and neighborhood impoverishment, averaged across the first 3 assessments (age 1.5, 2, and 3.5). Familial income was assessed by mother report. Neighborhood impoverishment was quantified by combining several block level variables from census data collected in 1990 (Shaw et al., 2012). Both were converted to Z-scores, and the mean of these standardized scores was used as the composite measure of early SES.

***Psychopathology.*** Internalizing and externalizing scores from the parent report form of the Child Behavior Checklist (Achenbach & Edelbrock, 1983) were averaged across ages 10-12 because these assessments precede the onset of cannabis use. Both variables were log transformed to account for positive skew in the data.

***Educational attainment.*** At age 22, participants reported the highest level of education they had completed on a 13-point scale (ranging from 1=below grade 9 to 13=completion of graduate degree.) Analyses examined educational attainment continuously.

***Occupational attainment.*** Participants’ employment status (currently employed/student vs. unemployed during the last year) and their self-reported job satisfaction (“How happy are you with this job?”; 1=very happy to 5=very unhappy) were assessed with the Revised Work Characteristics and Unemployment Measure {Conger, 1988}, which was administered to participants over the phone at age 23.

**Diffusion Tensor Imaging (DTI).**

***DTI acquisition.*** Two axial 2D DTI bipolar scans were acquired using identical parameters at both ages: time-to-repetition (TR)=8400 ms; time-to-echo (TE)=91ms; field of view=256x256; frequency=96; phase=96; 64 slices of 2mm thickness were acquired for a total scan time=9 min and 56 s. Diffusion-sensitizing gradient encoding was applied in 61 uniform angular directions with a diffusion weighting of b=1000 s/mm^2^. Seven reference images with no diffusion gradient (b=0) were also acquired.

***DTI preprocessing.*** Preprocessing was carried out using the Oxford Centre for Functional MRI of the Brain (FMRIB) Software Library (FSL) (35) using tract-based spatial statistics (TBSS) (36), including brain extraction, eddy current correction, and fitting a tensor model at each voxel. All subjects' FA data were eroded, end slices were removed to eliminate likely outliers, and nonlinear registration was used to align all FA images into a common space (37, 38). A mean FA image was then created and thinned to create a mean FA skeleton onto which each subject's aligned FA data was then projected. For mean (MD), axial (AD), and radial (RD) diffusivity, the mean image was registered to the FA skeleton (given that AD and RD are subcomponents of FA, AD/RD results are presented in Supplemental Materials). The Johns Hopkins University White Matter Tractography Atlas (39) was used to identify the right and left cingulum (cingulate gyrus) and ATR as regions of interests (ROIs), and mean FA, MD, AD, and RD values for each ROI were extracted to SPSS for further analysis. The cingulate gyrus ROI includes fibers that travel through coronal planes at both the middle of the splenium of the corpus callosum and the middle of the genu of the corpus callosum. The ATR ROI includes fibers that travel through coronal planes at the middle of the genu of the corpus callosum and the thalamus, excluding any fibers that cross the corpus callosum (39)

**Results**

**Cross-sectional Association between Adolescent Cannabis Use and ACC Connectivity at Age 20**

**Cingulum AD/RD.** Adolescent cannabis exposure did not have a significant effect on cingulum AD (*F*=2.57, *p*=.08) or RD (*F*=0.07, *p*=.93).

**ATR AD/RD.** A significant effect of adolescent cannabis exposure on ATR AD was observed (*F*=4.31, *p*=.014; see Figure 9). Consistent with the pattern observed for ATR FA, the moderate exposure group had higher AD than both other groups. Post-hoc tests demonstrated that the pairwise difference between the moderate and heavy exposure groups was statistically significant (*p*_corrected_=.012). This effect remained significant (*F*=4.28, *p*=.015) after controlling for alcohol exposure (*F*=6.4, *p*=.012), and post-hoc pairwise tests revealed a significant difference between the low/non-using group and the moderate use group (*p*_corrected_=.02). Adolescent cannabis exposure did not have a significant effect on ATR RD at age 20 (*F*=2.4, *p*=.09).

**Longitudinal Association between Extended Cannabis Use and Developing ACC Connectivity from age 20-22**

**Cingulum AD/RD change from age 20 to 22*.*** No significant effect of cumulative cannabis exposure or hemisphere was observed for change in cingulum AD or RD from age 20 to 22.

**ATR AD/RD change from age 20 to 22.** The pattern of change in ATR RD was consistent with the results for ATR FA: whereas the moderate exposure group displayed the largest *decrease* in ATR FA, they also displayed the largest *increase* in ATR RD from age 20 to 22. ANOVA results demonstrated a significant effect of cumulative cannabis exposure on change in ATR RD (*F*=3.28, *p*=.039; see Figure 11). Bonferroni-corrected post-hoc pairwise tests showed that the difference between the no/low exposure and moderate exposure groups was statistically significant (*p*_corrected_=.037). Change in ATR RD did not differ significantly based on hemisphere. No significant effect of cumulative cannabis exposure or hemisphere was observed for change in ATR AD from age 20 to 22.

Supplementary Table 1. Model Selection

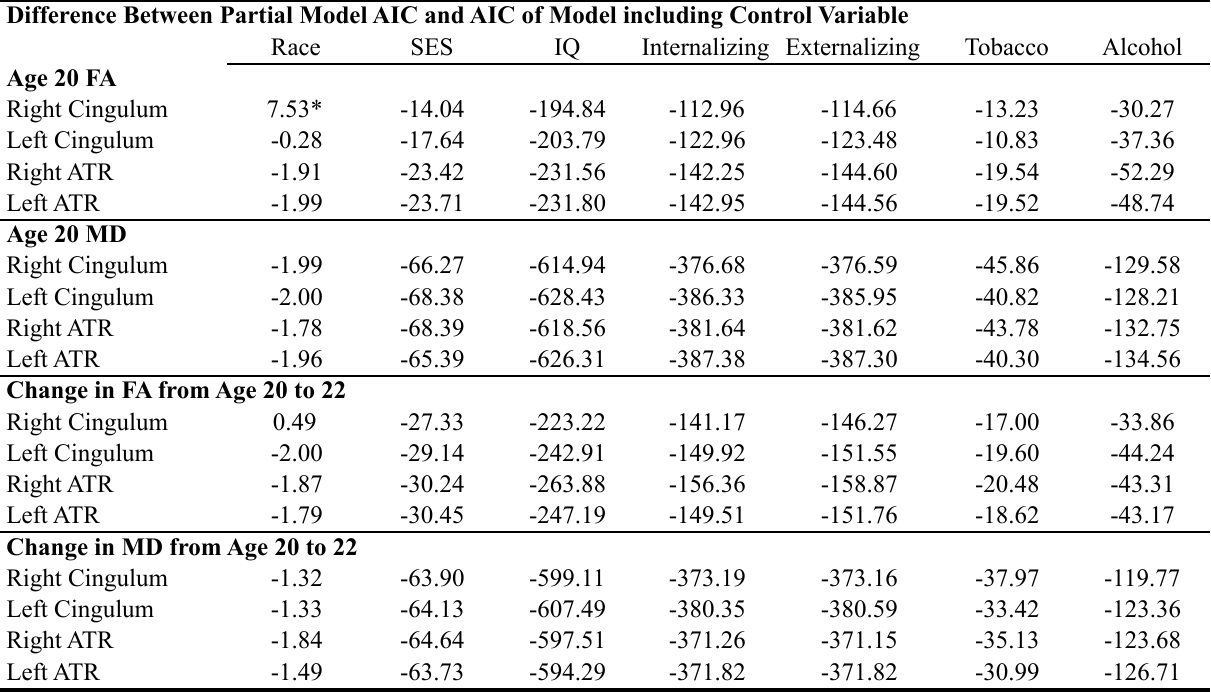

*Note.* Akaike Information Criterion (AIC) values were calculated for regression models predicting white matter microstructure (FA and MD of the right and left cingulum and ATR) based on adolescent cannabis exposure, controlling for head motion during DTI scanning. These were compared to AIC values for regression models including each of the potential covariates. Smaller AIC values reflect better regression model fit. Therefore, the AIC values from the models including our control variables were subtracted from the AIC value of the original model. Difference values greater than 2 reflect a substantial improvement in model fit. Race was the only control variable that improved the model fit for right cingulum FA at age 20. Therefore, race was included as a covariate in regression models predicting FA, AD, and RD at age 20.

Supplementary Table 2. Alcohol and Other Substance Use Characteristics

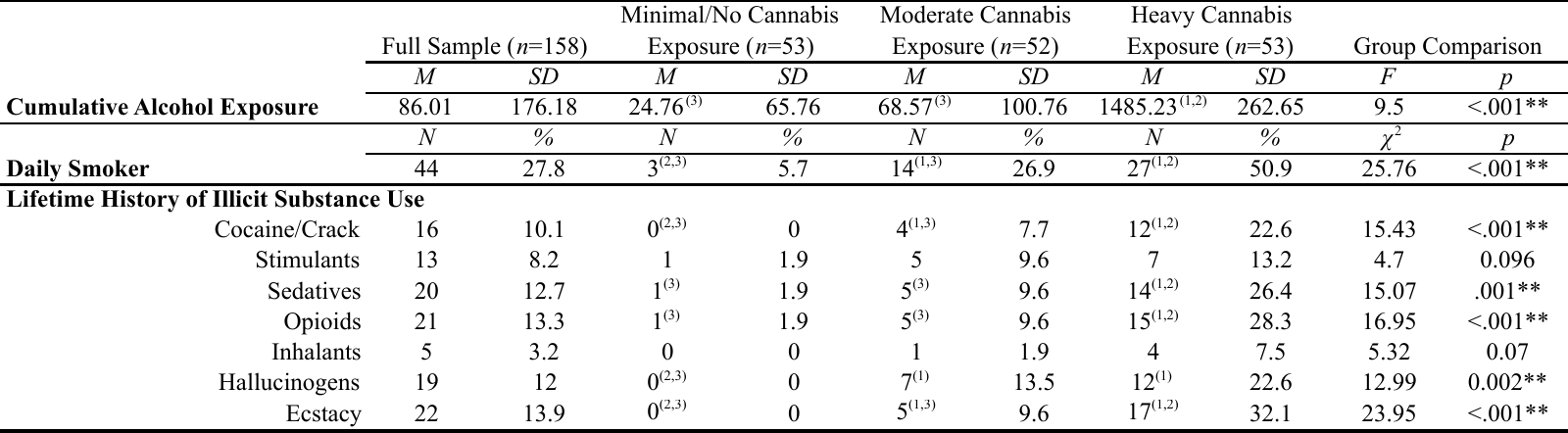

*Note*. **p*<.05 ***p*<.01. Superscript numbers in parentheses indicate which groups were significantly different from one another, based on pairwise Bonferroni-corrected post-hoc testing or pairwise *χ*2 tests, as applicable (1=Minimal/No Cannabis Exposure Group, 2=Moderate Cannabis Exposure Group, 3=Heavy Cannabis Exposure Group). Cumulative alcohol exposure reflects the sum of participants’ annual quantity of alcohol use (average days/month and average drinks/occasion were multiplied in order to obtain a measure of overall quantity of alcohol exposure for each year). Lifetime history of illicit substance use was assessed using the Lifetime History of Drug Use and Drug Consumption (LHDU) semi-structured interview; positive lifetime history was determined based by consensus from age 20 and age 22 study visits.

Supplementary Table 3. Cross-sectional Association between Adolescent Cannabis Use and ACC Connectivity at Age 20, controlling for Cumulative Alcohol Exposure

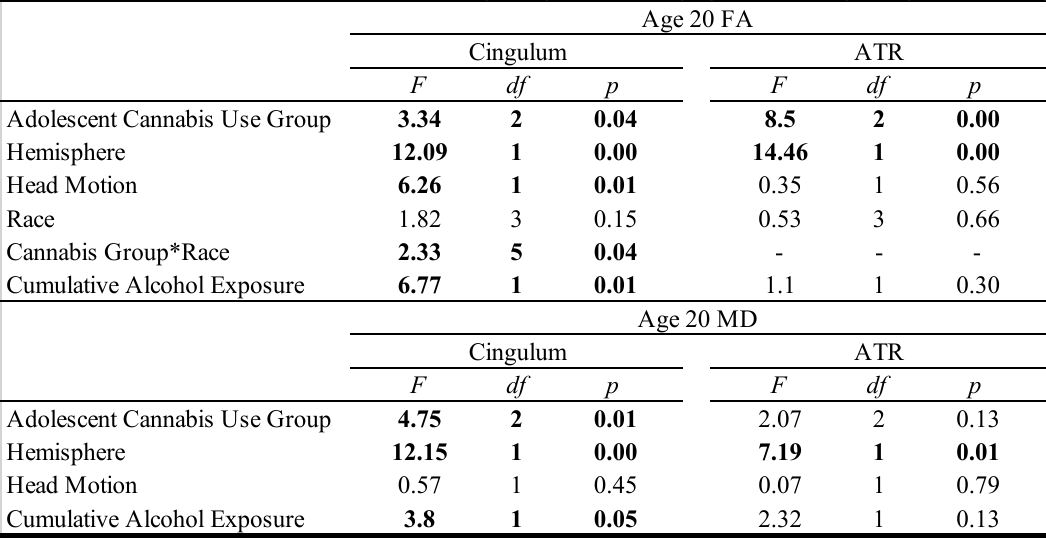

*Note*. Each quadrant represents one ANCOVA and significant effects (*p*<.05) are bolded. FA=fractional anisotropy, ATR=anterior thalamic radiations, *df*=degrees of freedom.

Supplementary Table 4. Longitudinal Association between Extended Cannabis Use and Developing ACC Connectivity from age 20-22, controlling for Cumulative Alcohol Exposure

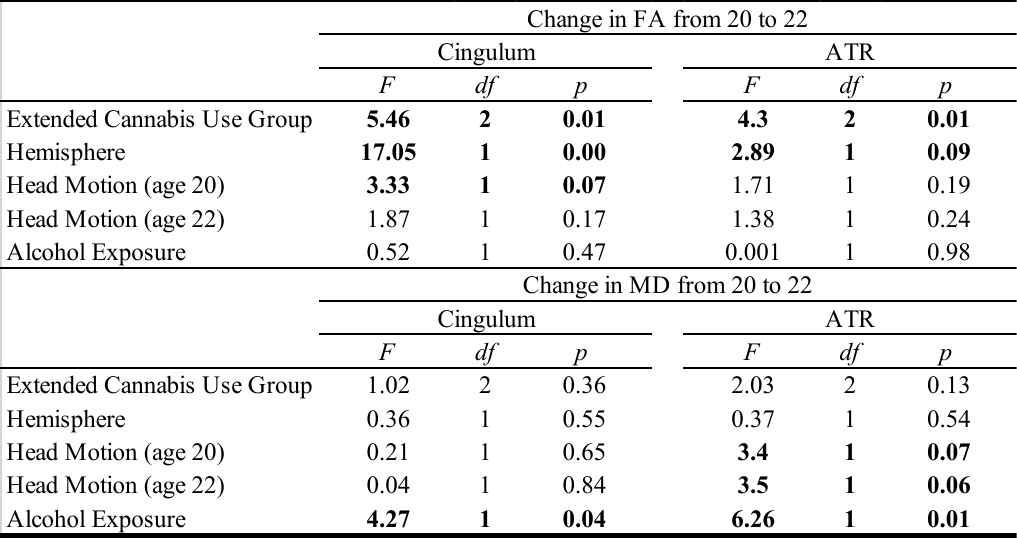

*Note*. Each quadrant represents one ANCOVA and significant effects (*p*<.05) are bolded. Change in FA from 20 to 22 represents the difference score for FA from age 20 to 22, and change in MD from 20 to 22 represents the difference score for MD from age 20 to 22. FA=fractional anisotropy, ATR=anterior thalamic radiations, *df*=degrees of freedom.
